# Transcriptomic Analysis of Human Skin Wound Healing and Rejuvenation Following Ablative Fractional Laser Treatment

**DOI:** 10.1101/2021.07.26.453869

**Authors:** Joseph D. Sherrill, Deborah Finlay, Robert L. Binder, Michael K. Robinson, Xingtao Wei, Jay P. Tiesman, Michael J. Flagler, Jean M. Loftus, Alexa B. Kimball, Charles C. Bascom, Robert J. Isfort

## Abstract

Ablative fractional laser treatment is considered the gold standard for skin rejuvenation. In order to understand how fractional laser works to rejuvenate skin, we performed microarray profiling on skin biopsies to identify temporal and dose-response changes in gene expression following fractional laser treatment. The backs of 14 women were treated with ablative fractional laser (Fraxel®) and 4 mm punch biopsies were collected from an untreated site and at the treated sites 1, 3, 7, 14, 21 and 28 days after the single treatment. In addition, in order to understand the effect that multiple fractional laser treatments have on skin rejuvenation, several sites were treated sequentially with either 1, 2, 3, or 4 treatments (with 28 days between treatments) followed by the collection of 4 mm punch biopsies. RNA was extracted from the biopsies, analyzed using Affymetrix U219 chips and gene expression was compared between untreated and treated sites. We observed dramatic changes in gene expression as early as 1 day after fractional laser treatment with changes remaining elevated even after 1 month. Analysis of individual genes demonstrated significant and time related changes in inflammatory, epidermal, and dermal genes, with dermal genes linked to extracellular matrix formation changing at later time points following fractional laser treatment. When comparing the age-related changes in skin gene expression to those induced by fractional laser, it was observed that fractional laser treatment reverses many of the changes in the aging gene expression. Finally, multiple fractional laser treatments resulted in continued changes in gene expression, with many genes either differentially regulated or continuously upregulated with increasing number of treatments, indicating that maximal skin rejuvenation requires multiple fractional laser treatments. In conclusion, fractional laser treatment of skin activates several biological processes involved in wound healing and tissue regeneration, all of which significantly contribute to the rejuvenating effect of fractional laser treatment on aged skin.

## Introduction

Aging of epithelial tissues, such as skin, is a complex biological phenomenon involving the epidermis, dermis and sub-dermis. Analysis of skin aging using systems biology approaches such transcriptomics, RNASeq, proteomics, etc., have demonstrated that skin aging is a dynamic condition with epidermal, dermal and subdermal aging progressing through different processes (Lener et al., 2006; Makrantonaki et al., 2012; Sextius et al., 2015; Holzscheck et al., 2020; Yan et al., 2013; Blumenberg, 2019; Kuehne et al., 2017; Kimball et al., 2017; Xu et al., 2016; Sole-Boldo et al., 2020; Cho et al., 2018; Zou et al., 2020; Haustead et al., 2016). Multiple approaches including chemical peels, dermabrasion, laser treatment (e.g. Fraxel®), non-laser light treatments, needling/microneedling, photodynamic therapy, radiofrequency treatment (e.g. Thermage®), ultrasound (e.g. Ultherapy®) and combinations of the above have been utilized to rejuvenate aged skin by activating the innate wound repair processes in both the epidermis and dermis (Humphrey et al., 2016; Fathi et al., 2015; Orringer et al., 2008; Fernandes et al., 2008; Fitzpatrick et al., 1996; Helbig et al., 2010; Pryor et al., 2011; Goldman et al., 2010). Of these approaches, the laser-based wound/repair modality appears to have the best efficacy in terms of rejuvenating both the epidermis and dermis, with fractional laser treatment having the least negative patient experience (Brightman et al., 2009; Helbig and Paasch, 2011).

Lasers have been used since the early 1980s to deliver skin rejuvenation benefits. The first clinically used laser treatment was total ablative laser-based rejuvenation. This treatment results in significant dermal remodeling and epidermal regeneration that improves both skin texture and tone, however as a consequence of removal of most of the epidermis and significant amount of dermis, long healing times and significant side effect profiles are observed. Nonablative laser rejuvenation procedures were subsequently developed, and while these approaches reduced the damage to the epidermis, they had limited rejuvenation efficacy. Next, fractional ablative laser rejuvenation procedures were developed, using a grid pattern of separate individual micrometer laser shots to ablate only a fraction of the epidermis and dermis, resulting in dermal remodeling and improved texture and tone with much faster healing times and a significantly improved side effect profile (Brightman et al., 2009; Papadavid et al., 2003; Tierney et al., 2011; Cicchi et al., 2014). Of these three approaches, ablative laser rejuvenation is considered the best treatment in terms of efficacy for the treatment of photoaging, acne scars and rhytides, although it is associated with serious side effects including long healing times, scarring and delayed onset hypopigmentation. In contrast, because the laser does not injure the epidermis and just heats the dermis, non-ablative laser rejuvenation is very safe but not as effective as ablative lasers in skin rejuvenation. Finally, fractional ablative laser rejuvenation combines both the safety of non-ablative laser rejuvenation with the efficacy of ablative laser rejuvenation and has become the new gold standard for skin rejuvenation with long lasting rejuvenation effects in multiple ethnicities at multiple body sites (Brightman et al., 2009; Papadavid et al., 2003; Tierney et al., 2011; Helbig and Paasch, 2011; Tan et al., 2014; Ortiz et al., 2010; Sadick et al., 2009; Kohl et al., 2014; Grunewald et al., 2011).

Fractional laser skin rejuvenation works by creating microscopic thermal holes in a grid pattern through the epidermis and deep into the dermis, thus stimulating dermal and epidermal regeneration while leaving most (up to 95%) of the epidermis and dermis intact. This allows the laser-induced damage to be more quickly repaired (Hantash et al., 2007; Bedi et al., 2007; Rahman et al., 2009; Tierney et al., 2011; Allemann et al., 2010). The depth and width of the thermal damage and laser intensity can be altered to achieve maximal dermal rejuvenation, often with efficacy similar to ablative laser treatment, without lengthy re-epithelialization (2 days for fractional laser versus 2-3 weeks for ablative laser), fewer infections, fewer acneiform eruptions and greatly reduced erythema (Tierney et al., 2011; Cohen et al., 2017; Robati et al., 2017). Significant work has been performed to understand how ablative, non-ablative and fractional lasers work to rejuvenate skin. Histopathological studies have demonstrated that fractional laser treatment of skin results in columnar ablative zones surrounded by thermal coagulation zones containing denatured proteins in the epidermis and dermis. Epidermal repair occurs within 1-3 days, the epidermal/dermal junction is repaired by 7 days while dermal repair takes weeks to months (Bedi et al., 2006; Hantash et al., 2007; Laubach et al., 2006; Trelles et al., 2008). Also observed was an increase in epidermal thickness, an increase in dermal collagens and a decrease in dermal elastin (elastosis) (El-Domyati et al., 2013; El-Domyati et al., 2014). These observations have been confirmed using the non-invasive techniques of confocal microscopy and two-photon microscopy (Longo et al., 2013; Cicchi et al., 2014). Histochemical studies looking at specific proteins demonstrate an increase in collagens (COL-1, COL-3), tropoelastin, vimentin, cytokines (EGF, bFGF, PDGF, VEGF, TGF-β HSP47, HSP72), apoptosis markers (annexin-VII, Bcl-2, caspase-9), inflammatory cells (CD3+, CD20+, CD68+, CD31+), proteases (MMP1, MMP3, MMP9, neutrophil elastase) and a decrease in p53 (Prignano et al., 2009; Helbig et al., 2011; Xu et al., 2011; Prignano et al., 2012; Helbig et al., 2009; El-Domyati et al., 2007; Zheng et al., 2014). Gene expression studies (typically using qPCR evaluating a limited number of genes) demonstrate an increase in proteases (MMP1, MMP2, MMP3, MMP9, MMP10, MMP11, MMP13), protease inhibitors (TIMP), hyaluronic acid synthesis enzymes (HAS, HYAL), collagens (COL1A1, COL3A1), extracellular matrix components (ELN, EMILIN1), cytokines (IL-1β and Paasch, 2011; Reilly et al., 2010; Orringer et al., 2012; Starnes et al., 2012; Kislevitz et al., 2020). Many of the observations of the effects of fractional laser treatment on human skin have been confirmed using animal and human skin organotypic models with the advantage that more detailed time courses and larger, more numerous sampling can be performed (Bedi et al., 2007; Park et al., 2012; Jiang et al., 2012; Jiang et al., 2014; Natari et al., 2020; Qu et al., 2019; Amann et al., 2016; Schmitt et al., 2018; Huth et al., 2020; Schmitt et al., 2017; Marquardt et al., 2015; Nogita et al., 2017). These analyses have provided a more complete picture of fractional laser treatment; however, the human and/or *in vivo* relevance of these findings need to be confirmed by actual human studies. More recently, transcriptomic analyses have been performed using microarrays, gene chips and RNAseq approaches, and have identified several gene expression changes occurring following fractional laser treatment on human skin (Kim et al., 2013; Jiang et al., 2012; Kim et al., 2019). Because of the global nature of these approaches, many gene expression changes have been observed following fractional laser treatment, albeit the generation of a holistic understanding of how this treatment results in skin rejuvenation has been hampered by small and limited sampling. Thus, a study evaluating a larger base size of individuals over longer time periods, with frequent sampling using sensitive transcriptomics, is needed to better understand how fractional laser treatment rejuvenates skin. Here,we present our findings from a detailed time course transcriptomic analysis of in vivo fractional laser treated human skin with the goal of providing a holistic understanding of how fractional laser-induced wounding and healing rejuvenates aged human skin.

## Materials and methods

### Study Design

This study was approved by the Schulman Associates Institutional Review Board. All subjects provided written informed consent prior to undergoing any study procedures. Fourteen women aged 30-55 with Fitzpatrick skin type I-IV were enrolled in this study. Fractional laser treatments were performed using a Fraxel® laser set to 12mj and 250mtz with a 1 cm^2^ treatment area, with all treatments and biopsy sampling performed by licensed medical personnel. For this study, the 14 study participants were treated on the back at 9 different equally spaced sites between the neck, upper buttocks and the left/right edges of the back with a minimum distance of 5 cm between each treatment site (plus one untreated site), and all sites were tattooed for site recognition. Sixty minutes prior to Fraxel® laser treatment, the treatment site was dosed with a topical anesthetic (BLT Cream: 20% benzocaine, 6% lidocaine, 4% tetracaine) after which the hand-held Fraxel® laser was moved over the treatment site 8 times. The treatment groups were as follows: (1) single laser treatments at 1, 3, 7, 14, 21 and 28 days before biopsy and (2) multiple laser treatments at 28-day intervals with 1, 2, 3, and 4 treatments. Following the final treatment, a 4 mm full thickness biopsy was taken at each site (including a non-treated site), the biopsy site was sutured, and the biopsy sample was placed in a cryotube and flash frozen in liquid nitrogen. A total of 140 biopsy samples were generated for analysis. Study design, cohort description, and sample collection for the aged dermis study has been previously described (Kimball et al., 2018).

### Microarray profiling

Microarray profiling was performed on 140 untreated or laser-treated samples using the HG-U219 array (Affymetrix) as similarly described for the aged dermis samples (Kimball et al. 2018). Flash-frozen, bisected 4mm full-thickness skin punch biopsies were homogenized in TRIzol reagent (ThermoFisher, Waltham, MA) and total RNA was extracted according to the manufacturer’s protocol. RNA was further purified using RNEasy spin columns (QIAGEN, Hilden, Germany). Biotinylated cRNA libraries were synthesized from 500 ng of total RNA using the Affymetrix HT 3’ IVT Express kit (Affymetrix, Santa Clara, CA) and ta Beckman Biomek FXp Laboratory Automation Workstation (Beckman Coulter, Brea, CA). Biotinylated cRNA was subsequently fragmented by limited alkaline hydrolysis and then hybridized overnight to the Affymetrix Human Genome U219-96 GeneTitan array. Plates were scanned using the Affymetrix GeneTitan Instrument utilizing AGCC GeneTitan Instrument Control software version 4.2.0.1566. Image data was summarized using Affymetrix PLIER algorithm with quantile normalization. Microarray data from this study as well as the aged dermis study have been deposited in the Gene Expression Omnibus (GEO) under GSE168760 and GSE139300, respectively.

### Statistical Analyses

After rigorous quality check using methods and metrics described by Canales, et al, a linear mixed ANOVA model was fitted to the microarray data using SAS software PROC GLIMMIX, version 9.4 of the SAS System for Windows (Canales et al. 2006). Gene expression (in log scale) of each single treatment site was compared to that of the baseline (day 0) untreated site, and each of the multiple treatment sites was compared to the single treatment site at day 28. Subject, Affymetrix plate (samples from 14 subjects were processed on two plates with seven subjects per plate) and Site were class variables. Subject (nested within Plate) was the random effect in the model. No multiple comparison adjustment was used due to the exploratory nature of the study.

### Bioinformatic Analyses

Gene signature analyses were performed using GeneSpring version 14.8 (Agilent Technologies, Santa Clara, CA). In brief, low-expressed probe sets were removed a minimum expression threshold (>20th percentile across all samples within any one treatment group) and condensed to a single Entrez gene ID using the arithmetic mean. Log_2_ fold changes were calculated for the single treatment sites relative to untreated sites (Day 0) or, for the multiple treatment sites, relative to the single treatment site (Day 28). Hierarchical clustering of the log_2_ fold change values using Euclidean distance was performed to create the single treatment and multiple treatment dendrograms. Biological pathway enrichment was performed on the filtered single treatment signatures (*p* < 0.01, absolute fold change > 1.1) and multiple treatment signatures (*p* < 0.05) using Ingenuity Pathway Analysis version 01LJ07 (QIAGEN). Fisher’s exact t test was used to assess significantly associated pathways, and zLJscores were calculated to predict pathway activation states (activated pathways, zLJscore >0; inhibited pathways zLJscore <0).

## Results

In order to understand how fractional laser treatment works to rejuvenate human skin, an Affymetrix gene chip based transcriptomic analysis was performed on human skin biopsies from fractional laser treated skin at various time points after a single and multiple treatments. The experimental design was two-fold: (1) to understand the biological process that occurs during healing following one fractional laser treatment, 6 different sites on the backs of 14 individuals were treated with a single fractional laser treatment followed by biopsy sampling of those sites at 1, 3, 7, 14, 21 and 28 days after treatment; and (2) in order to understand if multiple treatments result in better skin rejuvenation, 3 different sites on the backs of the same 14 individuals received either 2, 3 or 4 fractional laser treatments at the same site (with the treatments 28 days apart) followed by biopsy sampling. All the single treatments were compared to an untreated biopsy sample, allowing the use of the same individual as their own control (see Materials and Methods for a detailed description). For single treatment sites, the number of differentially expressed probesets was highest 1 day after treatment and continued to drop throughout the 28- day treatment period (Table I). Interestingly, multiple treatments of the same site, spaced 28 days apart, showed an increase in differential gene expression when compared to a single treatment after 28 days (Table I). Investigating the nature of the gene expression changes with time after fractional laser treatment demonstrated both increases and decreases in gene expression with a gradual normalization over time (Figure 1A). Clustering analysis showed several interesting patterns including a sharp increase in gene expression at 1 day post treatment followed by a sharp decline between 3-7 days post treatment – this cluster contains a number of keratin genes, S100 family alarmin genes and metalloproteinase genes – indicative of early epidermal and dermal wound healing responses. Also observed was a sharp decrease in gene expression at 1 day post treatment followed by an increase in expression 3 days post treatment and normalization to background levels by 7 days post treatment of a wide variety of genes. Finally, a sharp decrease in gene expression at 1 day post treatment followed by a gradual increase in expression over the 28 day treatment period – this cluster contains a number of dermal extracellular matrix genes (Figure 1B-D).

**Figure 1.**
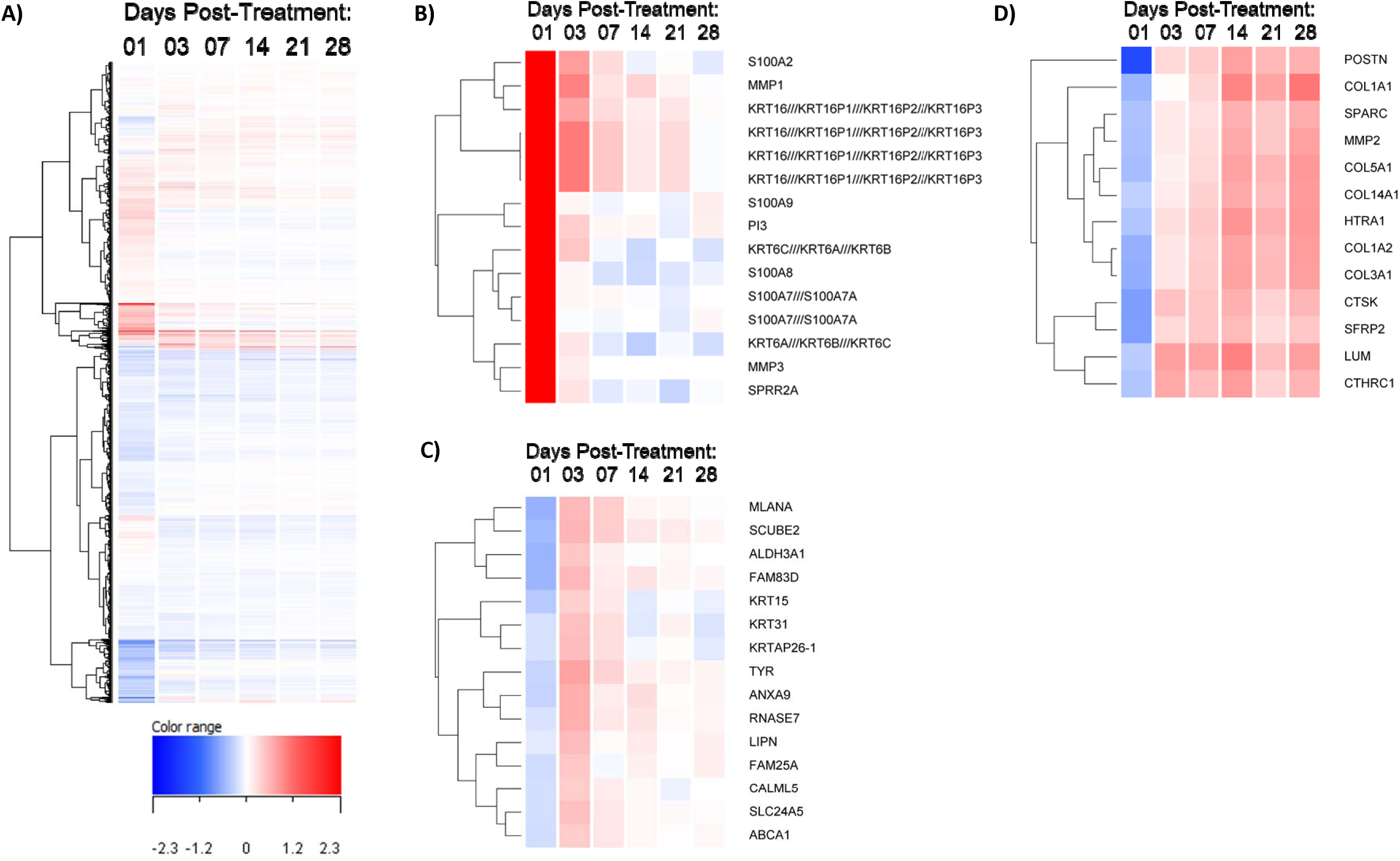
Kinetic effects of single laser resurfacing treatment on gene expression. **A)** Dendrogram displaying hierarchical clustering of genes by average log_2_ fold change that were significantly changed at any one time point compared to baseline untreated samples (n = 15,067 total). **B-D)** Dendrograms of select gene clusters from A) showing kinetic dysregulation from early (**B**, days 1-3), early-to-mid (**C**, days 3-7), and mid-to-late (**D**, days 14-28) time points following single laser resurfacing treatment.

**Table I.** Kinetic effects of laser resurfacing treatment(s) on gene expression in human skin.

| **Days post-treatment** | **Number of treatments** | **Comparison** | **No. differentially regulated probesets<sup>1</sup>** |
| --- | --- | --- | --- |
| 1 | 1 | vs Day 0<br>(untreated) | 24,358 |
| 3 | 1 |  | 16,754 |
| 7 | 1 |  | 11,473 |
| 14 | 1 |  | 10,693 |
| 21 | 1 |  | 5,456 |
| 28 | 1 |  | 6,555 |
| 56 | 2 | vs Day 28<br>(1 treatment) | 4,750 |
| 84 | 3 |  | 2,017 |
| 112 | 4 |  | 4,211 |

<sup>1</sup> Number of statistically significant probe sets (ANOVA $p$ -value < 0.05) expressed above the 30th percentile in $\geq 50\%$ samples in a treatment group

### Skin wound healing involves inflammatory, epidermal and dermal processes

A bioinformatic analysis of the top differentially regulated genes demonstrated a collection of activated inflammation related pathways was differentially regulated as early as 1 day after fractional laser treatment with most inflammatory processes normalized by 14 days post treatment (Table II). Epidermal related biological pathways involving proliferation and migration were similarly activated 1 day after treatment with many of the processes normalized by 7 days post treatment (Table III). Interestingly, in addition to proliferation and movement of epidermal cells, an epidermal-mesenchymal transition pathway was activated 3 to 7 days post treatment (Table III), which returned to baseline levels by day 14. This suggests the epidermis as a potential source for the mesenchymal cells participating in tissue repair at this early stage. Evaluation of dermal related biological pathways were complex with some processes initiating 1 day after fractional laser treatment such as proliferation and differentiation of connective tissue cells while other processes beginning much later such as the formation and maturation of collagen fibrils (Table IV). An analysis of the top 20 molecular functions 28 days post treatment supports the observations that cell death and removal, cell proliferation/movement and extracellular matrix maturation are the key processes during the wound healing process (Table V). A more granular display of the top 20 molecular functions over time and with multiple treatments is shown in Supplementary Tables 1-9.

**Table II.** Stress and immune response biological pathways associated with laser resurfacing kinetics.

| Annotation | z-score<sup>1</sup> |  |  |  |  |  | -log<sub>10</sub> p-value<sup>2</sup> |  |  |  |  |  |
| --- | --- | --- | --- | --- | --- | --- | --- | --- | --- | --- | --- | --- |
|  | Days Post-Treatment |  |  |  |  |  | Days Post-Treatment |  |  |  |  |  |
|  | 01 | 03 | 07 | 14 | 21 | 28 | 01 | 03 | 07 | 14 | 21 | 28 |
| Activation of leukocytes | N/A | 3.59 | 3.96 | 4.85 | N/A | 2.45 | 0.00 | 8.22 | 8.22 | 23.62 | 0.00 | 3.92 |
| Adhesion of immune cells | N/A | 5.04 | 4.27 | 4.81 | N/A | N/A | 0.00 | 8.18 | 9.59 | 21.44 | 0.00 | 0.00 |
| Antimicrobial response | N/A | N/A | 2.21 | 2.71 | 1.55 | 1.73 | 0.00 | 0.00 | 10.48 | 14.79 | 12.35 | 4.22 |
| Cell movement of lymphocytes | N/A | N/A | 4.46 | 4.34 | N/A | N/A | 0.00 | 0.00 | 7.62 | 18.89 | 0.00 | 0.00 |
| Cell movement of neutrophils | N/A | 3.32 | 2.69 | 3.62 | N/A | N/A | 0.00 | 12.13 | 10.66 | 15.95 | 0.00 | 0.00 |
| Degranulation of granulocytes | 2.11 | 3.06 | N/A | 2.54 | N/A | N/A | 18.10 | 14.10 | 0.00 | 14.25 | 0.00 | 0.00 |
| DNA damage | -1.72 | N/A | N/A | N/A | N/A | N/A | 6.83 | 0.00 | 0.00 | 0.00 | 0.00 | 0.00 |
| Endocytosis | 3.04 | 3.71 | 3.02 | 3.34 | N/A | N/A | 10.13 | 9.86 | 10.50 | 11.32 | 0.00 | 0.00 |
| Engulfment of cells | 2.74 | 3.79 | 3.42 | 3.54 | N/A | N/A | 9.71 | 8.41 | 9.84 | 11.34 | 0.00 | 0.00 |
| Immune response of cells | 1.76 | 3.83 | 3.87 | 4.95 | N/A | N/A | 9.45 | 10.96 | 7.40 | 13.65 | 0.00 | 0.00 |
| Metabolism of reactive oxygen species | 3.71 | 1.84 | N/A | 3.10 | N/A | N/A | 7.53 | 9.38 | 0.00 | 8.68 | 0.00 | 0.00 |
| Proliferation of immune cells | 1.62 | 2.06 | 2.96 | 1.74 | 1.86 | 1.37 | 8.57 | 11.22 | 9.27 | 26.51 | 3.57 | 3.49 |
| Recruitment of granulocytes | N/A | N/A | N/A | 3.02 | N/A | N/A | 0.00 | 0.00 | 0.00 | 8.76 | 0.00 | 0.00 |
| Response of phagocytes | N/A | 3.17 | 2.88 | 4.30 | N/A | N/A | 0.00 | 8.64 | 7.93 | 16.92 | 0.00 | 0.00 |
| Synthesis of reactive oxygen species | 3.66 | 1.87 | N/A | 3.10 | N/A | N/A | 8.18 | 9.23 | 0.00 | 8.35 | 0.00 | 0.00 |
| T cell migration | N/A | N/A | N/A | 5.39 | N/A | N/A | 0.00 | 0.00 | 0.00 | 13.09 | 0.00 | 0.00 |
| T cell response | N/A | N/A | N/A | 3.27 | N/A | N/A | 0.00 | 0.00 | 0.00 | 8.11 | 0.00 | 0.00 |

<sup>1</sup> Activation z-score (see Methods); n/a, activation z-score not calculated
<sup>2</sup> Fisher's exact *p*-value

**Table III.** Epidermal-related biological pathways associated with laser resurfacing kinetics.

|  |  | z-score<sup>1</sup> |  |  |  |  |  | -log10 p-value<sup>2</sup> |  |  |  |  |  |
| --- | --- | --- | --- | --- | --- | --- | --- | --- | --- | --- | --- | --- | --- |
|  |  | Days Post-Treatment |  |  |  |  |  | Days Post-Treatment |  |  |  |  |  |
|  | Annotation | 01 | 03 | 07 | 14 | 21 | 28 | 01 | 03 | 07 | 14 | 21 | 28 |
| **Growth & Differentiation** | Cell movement of epithelial cells | 1.39 | 0.44 | -0.44 | 0.54 | 1.51 | 0.84 | 7.32 | 7.81 | 8.11 | 7.84 | 3.38 | 3.97 |
|  | Epithelial-mesenchymal transition | N/A | 1.35 | 1.89 | N/A | N/A | N/A | 0.00 | 8.99 | 8.61 | 0.00 | 0.00 | 0.00 |
|  | Proliferation of epithelial cells | 2.18 | -1.28 | 0.56 | N/A | N/A | N/A | 8.26 | 10.88 | 7.54 | 0.00 | 0.00 | 0.00 |
|  | Necrosis of epithelial tissue | 0.06 | 0.56 | N/A | N/A | N/A | N/A | 13.48 | 14.79 | 0.00 | 0.00 | 0.00 | 0.00 |

<sup>1</sup> Activation z-score (see Methods); n/a, activation z-score not calculated
<sup>2</sup> Fisher's exact *p*-value

**Table IV.** Dermal-related biological pathways associated with laser resurfacing kinetics.

|  |  | z-score<sup>1</sup> |  |  |  |  |  | -log10 p-value<sup>2</sup> |  |  |  |  |  |
| --- | --- | --- | --- | --- | --- | --- | --- | --- | --- | --- | --- | --- | --- |
|  |  | Days Post-Treatment |  |  |  |  |  | Days Post-Treatment |  |  |  |  |  |
|  | Annotation | 01 | 03 | 07 | 14 | 21 | 28 | 01 | 03 | 07 | 14 | 21 | 28 |
| **Growth & Differentiation** | Angiogenesis | 1.24 | 2.14 | 3.28 | 2.36 | 0.94 | 0.07 | 20.35 | 22.51 | 16.15 | 23.10 | 9.49 | 8.08 |
|  | Cell death of fibroblasts | -0.64 | N/A | N/A | N/A | N/A | N/A | 6.70 | 0.00 | 0.00 | 0.00 | 0.00 | 0.00 |
|  | Cell movement of connective tissue cells | 0.42 | 1.99 | 0.57 | N/A | N/A | N/A | 10.09 | 8.88 | 7.49 | 0.00 | 0.00 | 0.00 |
|  | Cell movement of dermal cells | N/A | N/A | N/A | N/A | N/A | 1.41 | 0.00 | 0.00 | 0.00 | 0.00 | 0.00 | 3.65 |
|  | Cell movement of vascular smooth muscle cells | N/A | N/A | N/A | N/A | 1.54 | 2.76 | 0.00 | 0.00 | 0.00 | 0.00 | 4.59 | 4.32 |
|  | Differentiation of connective tissue cells | N/A | 0.82 | 0.02 | 0.56 | 0.79 | 0.41 | 0.00 | 13.23 | 10.00 | 12.88 | 10.50 | 4.96 |
|  | Growth of connective tissue | 0.99 | 1.11 | 2.05 | 0.35 | 0.44 | 0.33 | 13.71 | 15.73 | 17.89 | 13.03 | 4.29 | 4.93 |
|  | Proliferation of connective tissue cells | 1.43 | 1.33 | 1.98 | 0.41 | 0.26 | -0.04 | 14.33 | 14.57 | 16.63 | 10.83 | 4.61 | 4.87 |
|  | Quantity of connective tissue | N/A | N/A | N/A | N/A | 0.16 | 2.17 | 0.00 | 0.00 | 0.00 | 0.00 | 3.57 | 3.87 |
|  | Vasculogenesis | 0.41 | 1.58 | 2.83 | 1.91 | 0.66 | -0.35 | 16.85 | 17.46 | 9.48 | 16.18 | 6.47 | 5.28 |
| **Matrix Development** | Adhesion of connective tissue cells | N/A | N/A | N/A | N/A | N/A | 1.87 | 0.00 | 0.00 | 0.00 | 0.00 | 0.00 | 3.30 |
|  | Attachment of cells | N/A | N/A | N/A | 1.53 | 1.04 | 1.39 | 0.00 | 0.00 | 0.00 | 10.67 | 3.28 | 5.53 |
|  | Cell spreading of fibroblasts | N/A | N/A | N/A | N/A | 0.90 | N/A | 0.00 | 0.00 | 0.00 | 0.00 | 5.02 | 0.00 |
|  | Fibrogenesis | -0.27 | 1.00 | 0.92 | 2.05 | 2.08 | 2.07 | 14.62 | 16.76 | 17.43 | 11.87 | 5.36 | 6.73 |
|  | Formation of collagen fibrils | N/A | N/A | N/A | N/A | 1.07 | 1.07 | 0.00 | 0.00 | 0.00 | 0.00 | 4.19 | 4.58 |
|  | Tensile strength of skin | N/A | N/A | N/A | N/A | 2.20 | 2.60 | 0.00 | 0.00 | 0.00 | 0.00 | 5.54 | 7.90 |

<sup>1</sup> Activation z-score (see Methods); n/a, activation z-score not calculated
<sup>2</sup> Fisher's exact *p*-value

**Table V.** Top 20 molecular functions and system development process annotations associated with day 28 post-laser resurfacing treatment (by Fisher’s p-value)

| Annotation | z-score<sup>1</sup> | p-Value<sup>2</sup> | # Molecules<sup>3</sup> |
| --- | --- | --- | --- |
| Cell movement | 2.38 | 2.07E-18 | 242 |
| Organization of extracellular matrix | n/a | 2.41E-17 | 48 |
| Organization of collagen fibrils | n/a | 1.83E-14 | 19 |
| Migration of cells | 2.25 | 7.78E-14 | 208 |
| Necrosis | -0.18 | 9.00E-13 | 250 |
| Apoptosis | -0.04 | 2.60E-12 | 244 |
| Morphology of collagen fibrils | n/a | 6.14E-12 | 11 |
| Morphology of fibrils | n/a | 8.09E-11 | 12 |
| Metabolism of protein | 1.13 | 4.20E-10 | 125 |
| Abnormal morphology of collagen fibrils | n/a | 1.47E-09 | 9 |
| Organization of filaments | 1.90 | 1.79E-09 | 35 |
| Angiogenesis | 0.07 | 8.33E-09 | 106 |
| Tensile strength of skin | 2.60 | 1.27E-08 | 9 |
| Development of vasculature | 0.07 | 2.20E-08 | 113 |
| Initiation of translation of protein | n/a | 2.63E-08 | 23 |
| Cell death of tumor cell lines | -0.74 | 2.87E-08 | 152 |
| Quantity of cells | 3.49 | 7.38E-08 | 168 |
| Formation of filaments | 2.19 | 1.61E-07 | 54 |
| Fibrogenesis | 2.07 | 1.87E-07 | 58 |
| Invasion of cells | 0.73 | 4.32E-07 | 101 |

<sup>1</sup> Activation z-score (see Methods); n/a, activation z-score not calculated
<sup>2</sup> Fisher's exact *p*-value
<sup>3</sup> Number of genes differentially regulated at day 28 within the annotated pathway

An evaluation of the canonical signaling pathways associated with fractional laser treatment demonstrated that apoptosis/proliferation, stress/inflammation and cell adhesion were the key pathway groupings activated (Table VI). In these 3 key pathway groupings, individual pathways demonstrated unique kinetics with peak activation of some pathways such as remodeling of epithelial adherens junctions, ephrin receptor signaling, cell cycle control, and inflammation (IL-6 and IL-8 signaling) occurring at early time points (days 1 – 7). Other pathways including activated interferon signaling and negative inhibition of MMP signaling remained elevated throughout the time points. Finally, the HOTAIR regulatory pathway remained active the entire 28 days of observation (Table VI and Figure 2).

**Figure 2.**
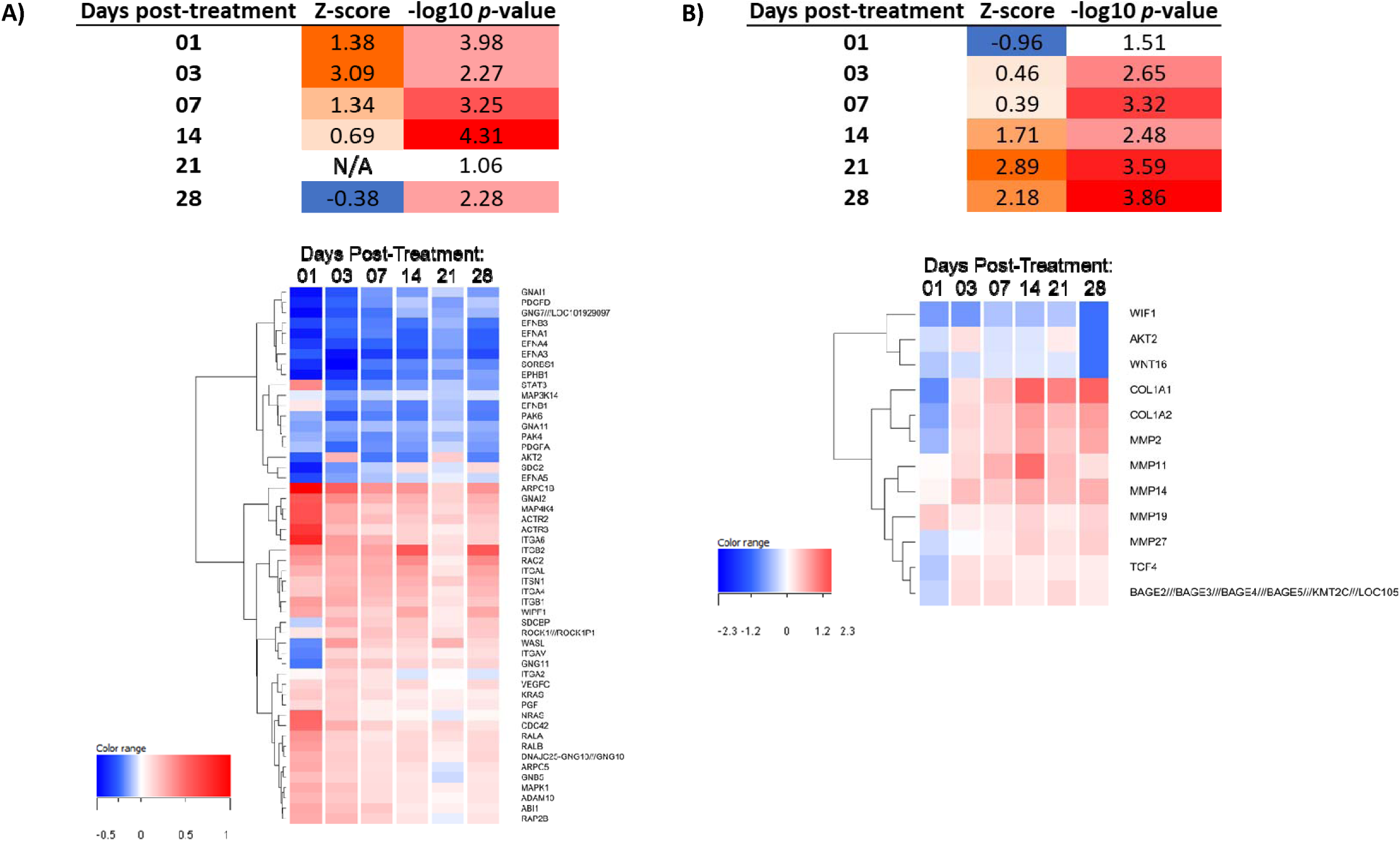
Single laser resurfacing treatment stimulates signaling pathways associated with wound repair. A) Activation of ephrin receptor signaling at early (days 1-3) time points following single laser resurfacing treatment. Dendrogram shows clustering of genes (n = 52) involved in the ephrin receptor signaling pathway by average log_2_ fold change compared to baseline untreated samples. B) Activation of the HOTAIR regulatory pathway at mid-to-late (days 14-28) time points following single laser resurfacing treatment. Dendrogram shows clustering of genes (n = 12) involved in the HOTAIR regulatory pathway by average log_2_ fold change compared to baseline untreated samples. Color scales for z-scores indicate predicted low/inhibition (blue) to high/activation (orange) states for each pathway at a given time point. Color scale for –log_10_ p-value reflect low (white) to high (red) significance for each pathway at a given time point.

**Table VI.** Canonical signaling pathways associated with laser resurfacing kinetics.

|  | Annotation | z-score<sup>1</sup> |  |  |  |  |  | -log<sub>10</sub> p-value<sup>2</sup> |  |  |  |  |  |
| --- | --- | --- | --- | --- | --- | --- | --- | --- | --- | --- | --- | --- | --- |
|  |  | Days Post-Treatment |  |  |  |  |  | Days Post-Treatment |  |  |  |  |  |
|  |  | 01 | 03 | 07 | 14 | 21 | 28 | 01 | 03 | 07 | 14 | 21 | 28 |
| **Apoptosis & Proliferation** | 14-3-3-mediated Signaling | 2.45 | 2.48 | 0.71 | 0.00 | N/A | N/A | 2.74 | 2.77 | 0.00 | 0.00 | 0.00 | 0.00 |
|  | Cell Cycle Control of Chromosomal Replication | 3.55 | 3.71 | 2.53 | N/A | N/A | N/A | 3.17 | 2.78 | 1.48 | 0.00 | 0.00 | 0.00 |
|  | Cyclins and Cell Cycle Regulation | 1.72 | 2.45 | 2.00 | -0.38 | N/A | N/A | 2.54 | 4.13 | 5.02 | 0.61 | 0.23 | 0.00 |
|  | HOTAIR Regulatory Pathway | -0.96 | 0.46 | 0.39 | 1.71 | 2.89 | 2.18 | 1.51 | 2.65 | 3.32 | 2.48 | 3.59 | 3.86 |
|  | HGF Signaling | 0.27 | 0.56 | -0.28 | -1.00 | N/A | -2.00 | 3.22 | 2.92 | 1.85 | 1.33 | 0.48 | 0.64 |
|  | Kinetochores Metaphase Signaling Pathway | 1.76 | 4.13 | 3.29 | 1.67 | N/A | N/A | 1.21 | 14.39 | 11.91 | 0.47 | 0.36 | 0.00 |
| **Stress & Inflammation** | EIF2 Signaling | -2.35 | -1.23 | -2.40 | -2.52 | 0.00 | -1.90 | 17.24 | 5.42 | 5.71 | 5.03 | 0.42 | 5.32 |
|  | ERK/MAPK Signaling | -0.11 | -0.43 | -0.39 | 0.00 | N/A | 0.00 | 3.05 | 2.47 | 2.46 | 1.46 | 0.27 | 0.00 |
|  | IL-6 Signaling | 1.21 | 1.41 | 0.54 | 0.28 | N/A | 0.71 | 3.13 | 1.30 | 0.54 | 0.73 | 0.44 | 0.80 |
|  | IL-8 Signaling | 1.12 | 2.02 | 0.96 | 2.27 | 0.00 | 0.58 | 4.70 | 3.16 | 1.58 | 2.59 | 0.28 | 0.92 |
|  | Inhibition of ARE-Mediated mRNA Degradation Pathway | 1.00 | 3.02 | 0.28 | 1.27 | N/A | N/A | 2.57 | 0.70 | 0.48 | 0.00 | 0.00 | 0.00 |
|  | Interferon Signaling | 3.58 | 3.64 | 2.89 | 3.46 | 2.83 | 2.45 | 2.02 | 4.30 | 4.93 | 5.36 | 6.84 | 3.21 |
|  | NF-κB Signaling | 0.97 | 2.38 | 0.66 | 1.15 | N/A | 1.00 | 2.48 | 2.93 | 0.93 | 1.03 | 0.00 | 0.54 |
| **Cell Adhesion** | Actin Cytoskeleton Signaling | 1.99 | 1.36 | -0.35 | -0.43 | N/A | -0.71 | 2.21 | 3.38 | 4.11 | 1.87 | 0.00 | 0.35 |
|  | Ephrin Receptor Signaling | 1.38 | 3.09 | 1.34 | 0.69 | N/A | -0.38 | 3.98 | 2.27 | 3.25 | 4.31 | 1.06 | 2.28 |
|  | Inhibition of Matrix Metalloproteases | -0.94 | 0.00 | -0.38 | -0.91 | -1.89 | -1.00 | 1.35 | 0.93 | 1.23 | 4.32 | 4.50 | 4.70 |
|  | Integrin Signaling | 1.55 | 2.18 | 0.85 | 1.89 | 0.00 | 0.28 | 6.05 | 2.43 | 4.15 | 2.59 | 0.63 | 1.31 |
|  | Rac Signaling | 2.21 | 1.52 | 0.47 | -0.54 | N/A | -2.45 | 3.97 | 4.13 | 3.57 | 2.20 | 0.25 | 0.89 |
|  | Remodeling of Epithelial Adherens Junctions | 3.30 | 1.00 | 0.00 | N/A | N/A | N/A | 3.95 | 1.28 | 1.07 | 0.00 | 0.00 | 0.00 |

<sup>1</sup> Activation z-score (see Methods); n/a, activation z-score not calculated
<sup>2</sup> Fisher's exact *p*-value

Fractional laser treatment is very effective in reversing skin aging changes. Thus, the transcriptional changes induced by fractional laser resurfacing were compared with the gene expression changes that occur during facial epidermis and dermis aging. The overlap of genes in common between the fractional laser treatment differentially expressed genes at 28 days post treatment and the differentially regulated genes in 50 versus 20-year-old epidermis (Figure 3A and 3B) and the 50 versus 20-year-old dermis were compared (Figure 3C and 3D). A large number of genes in common was observed withseveral genes being expressed in opposite directions in the fractional laser samples compared to the 50 versus 20-year-old epidermis and dermis samples(Figure 2B and 2D). A closer analysis revealed several extracellular matrix-relatedgenes, in particular collagen genes, were the most highly upregulated with fractional laser treatment at day 28 yet showed the greatest decrease in expression in aged versus young skin (Figure 3B and 3D; Figure 4A-B). Notably, many of the genes shared across multiple pathways displayed an increasing expression pattern over time following fractional laser treatment while their expression was decreased in 50-year-old individuals (Figure 4C). This indicates one of the major effects of fractional laser treatment is the reversal of the loss of extracellular matrix with skin aging.

**Figure 3.**
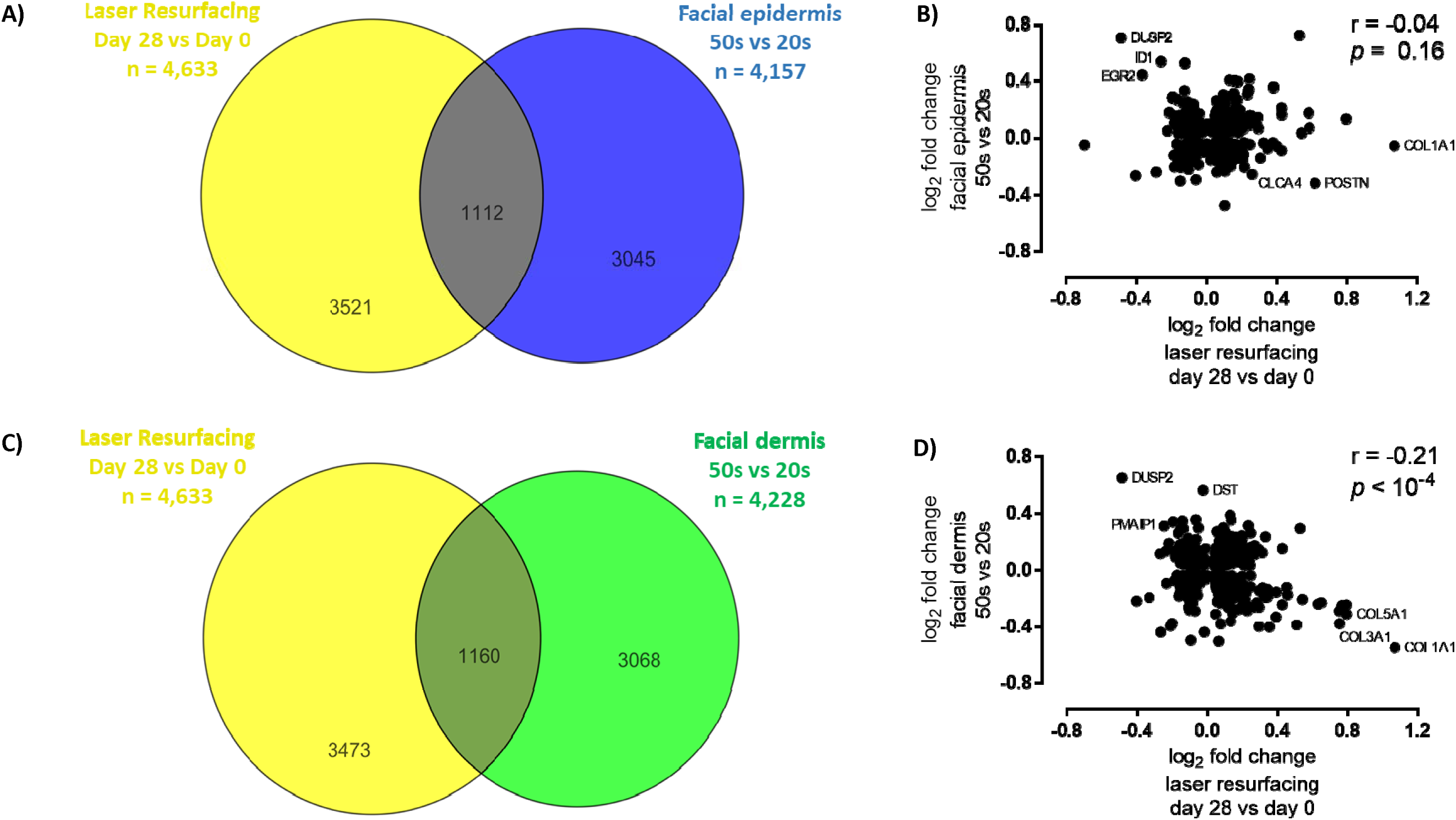
Laser resurfacing treatment reverses age-related gene expression changes. **A)** Venn diagram depicting the overlap in statistically significant genes altered at 28 days post-laser resurfacing and in facial epidermis between 50-year-olds and 20-year-olds. B) Log_2_ fold change (FC) plot and Pearson correlation for the overlapping genes from A). C) Venn diagram depicting the overlap in statistically significant genes in the laser resurfacing single-treatment kinetic signature and in facial epidermis between 50-year-olds and 20-year-olds. D) Log_2_ fold change (FC) plot and Pearson correlation for the overlapping genes from C).

**Figure 4.**
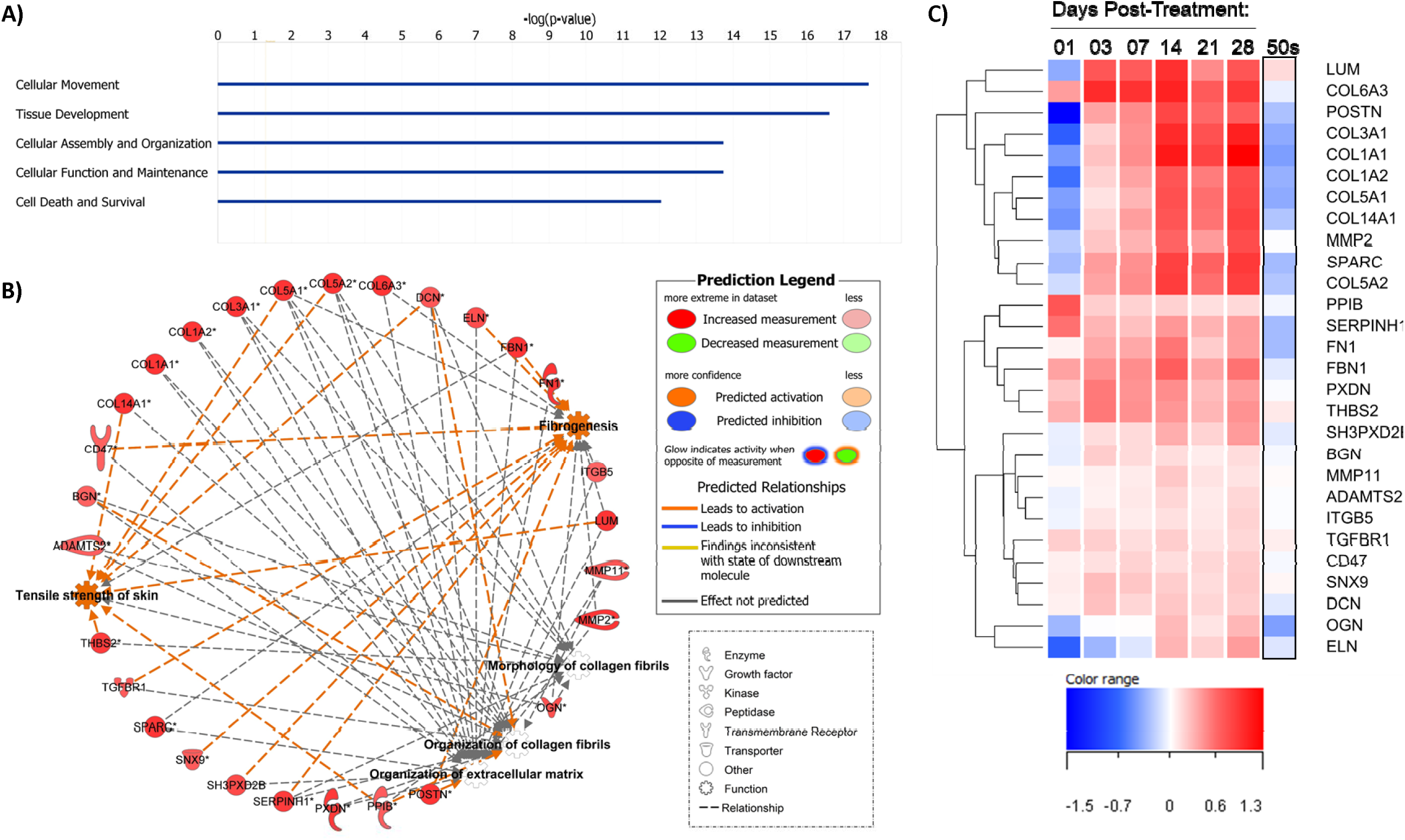
Activation of multiple collagen-related pathways by day 28 post-treatment. **A)** Bar chart showing top 5 molecular functions and system development processes associated with the day 28 gene signature (by –log_10_ Fisher’s p-value). B) Network of shared genes across select collagen-related pathways associated with day 28 gene signature (from Table VI). C) Dendrogram displaying hierarchical clustering of shared collagen across single laser resurfacing treatments (vs. baseline untreated samples) and in facial dermis of 50-year-olds vs. 20-year-olds by log_2_ fold change.

Finally, based on the clinical observation that multiple fractional laser treatments result in the best skin rejuvenation outcome, the gene expression changes that occur following 2, 3 or 4 sequential fractional laser treatments were examined (Figure 5). When compared to a single fractional laser treatment, increased numbers of differential gene expression changes were observed following 2 treatments (Figure 5A), many of which were maintained following 3 and 4 treatments and some which were only observed following 3 treatments (Figure 5B and 5C). An evaluation of two specific genes illustrated that multiple treatments are required for maximal effect for COL1A1 expression (Figure 5D) and NR4A1 (Figure 5E), albeit with different trajectories.

**Figure 5.**
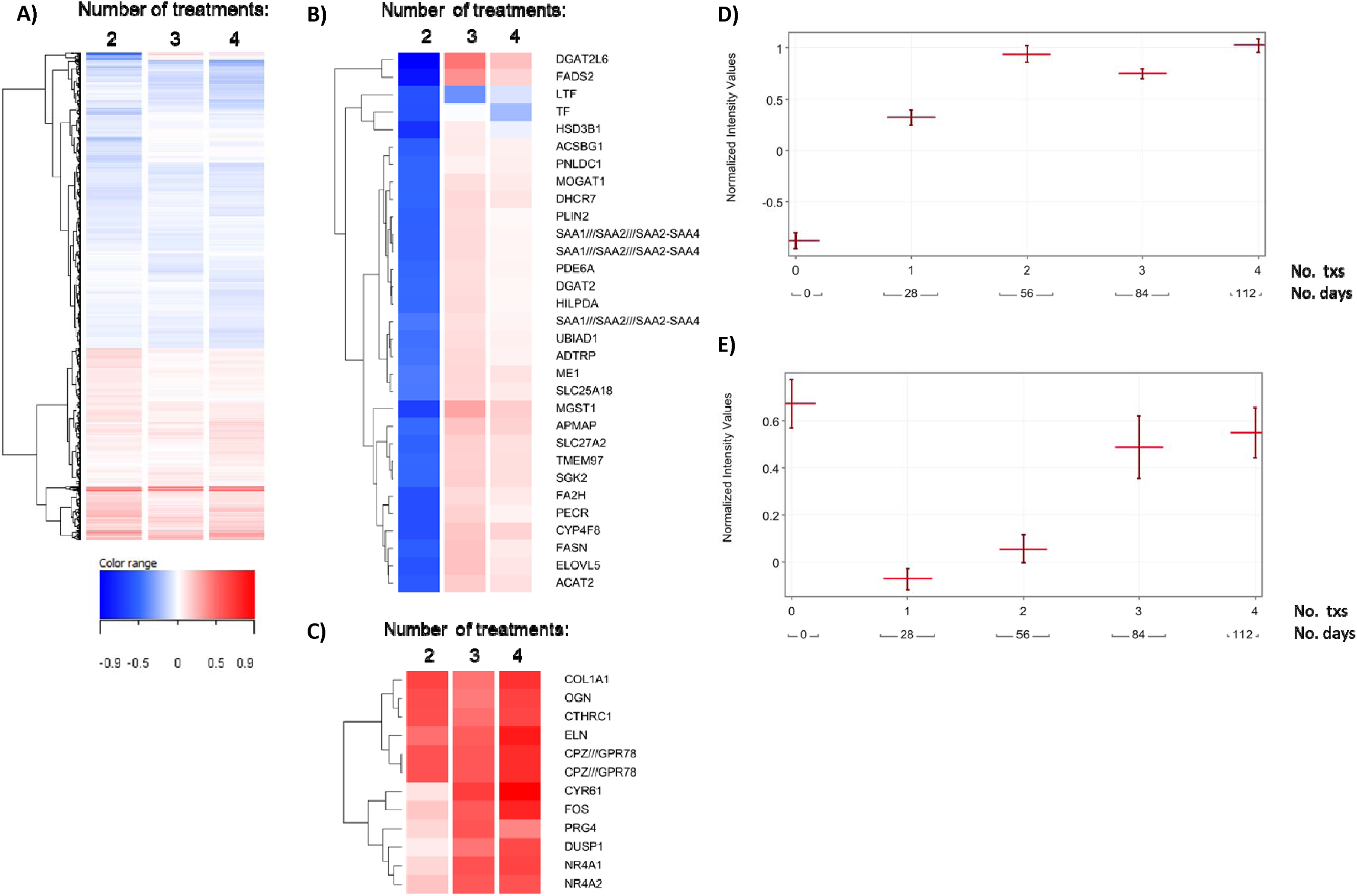
Effects of multiple laser resurfacing treatments over time on gene expression. **A)** Dendrogram displaying hierarchical clustering of genes by average log_2_ fold change that were significantly changed at any multiple-treatment sites compared to the single-treatment site (n = 5,795 total). B-C) Dendrograms of select gene clusters differentially regulated from 2-4 laser resurfacing treatments (B) or continuously upregulated with increasing number of treatments (C). D-E) Profile plots showing normalized expression values (±SEM) of COL1A1 (D) and NR4A1 (E) across the different number of treatments over time.

## Discussion

In this report, temporal transcriptomic analysis has been utilized to better understand how fractional laser treatment functions to rejuvenate aged skin. We demonstrate that fractional laser treatment induces many biological processes related to wound healing and tissue regeneration, in a highly orchestrated manner, resulting in rejuvenation of long-lived skin macromolecules like components of the extracellular matrix. These data greatly expand the previously published temporal analysis of the effects of laser treatment on skin by providing a much more detailed analysis of the changes in gene expression that occur (whole genome) compared to the approximately 20 genes previously analyzed (Helbig and Paasch, 2011; Reilly et al., 2010; Orringer et al, 2012, Starnes et al., 2012; Kislevitz et al., 2020). In addition, this study greatly extends previous transcriptomic analyses by including more sampling times, larger number of study participants and increased number of laser treatments (Kim et al, 2013; Jiang et al., 2012; Kim et al., 2019).

As part of the wound healing process following fractional laser-induced treatment, immediate gene expression changes were observed indicative of epidermal and dermal repair mediated by keratinocytes, fibroblasts and immune cells which were followed by longer term repair of the dermis mediated by fibroblasts and immune cells. Activation of keratinocytes as early as 1 day post fractional laser treatment is exemplified by the large changes in keratin gene expression and the activation of epithelial cell pathways (necrosis of epithelial tissues, proliferation and cell movement of epithelial cells) and is consistent with current knowledge concerning the speed of epidermal repair following fractional laser treatment (Bedi et al., 2006; Hantash et al., 2007; Laubach et al., 2006; Trelles et al., 2008). Two pathways important in the re-epithelization of a wound (ephrin receptor signaling pathway and HOTAIR regulatory pathway) were analyzed in more detail. The ephrin receptor signaling pathway, which is important in wound closure and healing (Nunan et al., 2015), is strongly activated as early as 1 day post treatment followed by a gradual decrease over the 28-day treatment period. The long non-coding RNA HOTAIR regulatory pathway was also activated following fractional laser treatment as would be required for proper wound healing (Shi et al., 2020). Additionally, the large changes in S100 gene expression at 1 day post fractional laser treatment, along with the rapid induction of gene expression of several inflammatory pathways, confirms a quick immune response to the damage and greatly expands previous histochemical study findings(Prignano et al., 2009; Helbig et al, 2011; Xu et al, 2011; Prignano et al, 2012; Helbig et al., 2009; El-Domyati et al., 2007; Zheng et al., 2014).

In the dermis, fibroblast mediated gene expression changes occurred immediately after treatment and continued throughout the four-month observation period. An interesting observation was the early (1 day after treatment) coordinated decrease in gene expression of fibroblast cell death and fibrogenesis pathways with an increase in gene expression of connective tissue growth, proliferation of connective tissue and cell movement of connective tissue pathways. These results indicate a complex fibroblast response to damage which appears to be focused on increasing fibroblast numbers and movement to damaged dermal areas. In contrast, later in the dermal wound healing process we observed an increase in the formation of collagen fibrils, tensile strength of skin, and attachment of cells pathways, all of which indicate a maturation of the wound healing response by synthesis and maturation of the extracellular matrix. The observation of these gene expression changes associated with activated tensile strength of skin, in particular, supports a direct relationship between the molecular changes induced by fractional laser treatment and the biomechanical properties of the skin. Activation of this pathway 28 days after treatment is consistent with the delayed onset of the skin appearance benefits following fractional laser treatment, and further supports that the late-stage dermal remodeling processes are crucial to the ultimate cosmetic benefits delivered by this gold standard anti-aging procedure. Together, the temporal analysis of gene expression changes following fractional laser wounding demonstrate a complex, yet orderly, repair process and provides significant insight into which biological processes are required for skin repair and rejuvenation.

Evaluation of the effects of multiple fractional laser treatment demonstrated that each treatment increases the skin rejuvenation response, best exemplified by the increase in COL1A1 and other extracellular matrix genes, with at least 2-3 treatments needed for maximal increase in gene expression. An evaluation of NR4A1, a nuclear receptor that functions as a transcriptional repressor of TGF β targeted genes whose expression is decreased during wound healing (Palumbo-Zerr et al., 2015), demonstrated at least 3 fractional laser treatments were required before NR4A1 levels increased to pretreatment levels. These observations fit well with the clinical experience in which maximal skin rejuvenation is observed following multiple fractional laser treatments (Allemann et al., 2010).

Multiple cosmetic procedures work to rejuvenate skin by activating the wound repair process (e.g., Fraxel® Thermage®, Ultherapy®), which among these approaches fractional laser is considered the gold standard treatment for aging related skin rejuvenation. The analysis of the gene expression changes observed following fractional laser treatment identified many biological and molecular pathways that are activated following treatment. These pathways were compared to the biological pathways that are modulated during skin aging using our previously described aging transcriptomics dataset (Kimball et al., 2017), in order to better understand how this treatment works for skin rejuvenation. A large number of genes in common between fractional laser treatment and aging of both the epidermis and dermis of human skin were observed, with treatment leading to reversal of many aging associated gene expression changes. These findings suggest that fractional laser treatment acts holistically across both epidermal and in dermal compartments to restore a more youthful skin structure. Many of the key biological processes affected by fractional laser treatment that are instrumental in skin rejuvenation are the processes involved in the regeneration of the dermal extracellular matrix. Reversal of the age associated loss of extracellular matrix gene expression (most notably collagen genes) was observed along with changes in gene expression indicative of significant remodeling of the extracellular matrix. These observations demonstrate that approaches to rejuvenate aged skin should ideally provide these two functions. Interestingly, several material-based approaches that achieve similar increases in extracellular matrix production to rejuvenate skin include peptides (such as matrikines) and combinations of peptides (Schagen 2017, Katayama 1991, Katayama 1993) and retinoids (Shao et al., 2017; Kong et al., 2016). Together these observations provide evidence that maximal skin rejuvenation requires a multi-faceted approach addressing both epidermal and dermal aging, with a key driver being the modulation of the long-lived dermal extracellular matrix. The transcriptome-wide effects of fractional laser induced skin repair and rejuvenation, analyzed in the present study with both a high degree of temporal precision and across multiple treatments, provides many new potential intervention targets to improve the structure, function and appearance of aging skin.

